# Metabolic properties of murine kidney mitochondria

**DOI:** 10.1101/2021.12.13.472400

**Authors:** Alexander V. Panov, Vladimir I. Mayorov, Sergey I. Dikalov

**Affiliations:** Scientific Centre for Family Health and Human Reproduction Problems, Irkutsk. Russian Federation; Department of Basic Medical Sciences, Mercer University School of Medicine, Macon, GA, Unated States of America; Division of Clinical Pharmacology, Vanderbilt University Medical Center, Nashville, TN Unated States of America

## Abstract

We show that mitochondria from the kidney of mice (MKM), rat brain (RBM), and heart (RHM) oxidize long-chain fatty acids at high rates in all metabolic states only in the presence of any other mitochondrial metabolites: succinate, glutamate, or pyruvate. All supporting substrates increased several folds the respiration rates in State 4 and State 3. The stimulations of the State 3 respiration with palmitoyl-carnitine + malate oxidation (100%) were: with succinate in MKM 340%, RBM 370%, and RHM 340%; with glutamate - MKM 200%, RBM 270%, and RHM 270%; and with pyruvate - MKM 150%, RBM 260%, and RHM 280%. The increases in O_2_ consumption in State 4 were due to increased leakage of electrons to produce superoxide radicals (O_2_^<u>•</u>^). Earlier, we have shown that the brain and heart mitochondria possess a strong intrinsic inhibition of succinate oxidation to prevent the excessive O_2_^<u>•</u>^ production at diminished functional loads. We show that kidney mitochondria lack the intrinsic inhibition of SDH. The new methodology to study β-oxidation of LCFAs opens the opportunity to study energy metabolism under normal and pathological conditions, particularly in the organs that utilize LCFAs as the main energy source.

## Introduction

All mammals have paired organs kidneys, designed to maintain the body’s water and ionic balances and purify the blood from contaminations. In humans, the kidneys in a day filtrate up to 180-200 liters of liquid containing ions and metabolites, which means that the whole body’s blood volume passes 36-40 times through the kidneys [1]. About 660 milliliters of blood plasma flow through both kidneys per minute. In the glomeruli, 125 ml of filtrate formed from this amount of plasma, which enters the lumen of the tubules, constituting approximately 19% of the plasma volume [2], and only a few milliliters become urine.

Glomerular filtration of the blood plasma with the primary urine formation does not directly require ATP or other types of metabolic energy. The driving force for glomerular filtration is the transcapillary difference in hydrostatic and oncotic pressures [2]. Meanwhile, almost all sodium ions and metabolites, such as glucose and amino acids, reabsorb back to the blood, which requires an enormous amount of energy. In addition to these excretory, ion, and osmolarity maintenance functions, kidneys also produce a large amount of glucose to maintain glucose homeostasis, together with the liver, during starvation [3]. Kidneys also have some hormonal and regulatory functions, requiring energy [4].

The central structural and functional units of a kidney are nephrons, which have the filtering part glomerulus and tubules where reabsorption occurs. Nephrons locate in the kidneys cortex [5]. Despite the delicate and complex molecular structures of the glomeruli and tubules containing many mitochondria [6,7], the metabolic functions of kidney mitochondria mostly remain an enigma. Physiologists a long time ago have established that fatty acids serve as a predominant substrate for the production of ATP [8,9]. Historically, physiologists much earlier than mitochondriologists have begun to understand the importance of the long-chain fatty acids in the energy supply of the organs’ functions. This notion applies not only to the kidneys but also to the brain and heart [10,11]. However, the isolated kidney mitochondria metabolic properties are much less studied than the mitochondria from other high energy-consuming organs, such as the heart and brain [12,13].

This paper presents new data on the metabolic properties of mouse kidney mitochondria obtained with the new methodology compared with the previously obtained data with the mitochondria of the rat heart [12] and the brain [13]. The central feature of the new approach to studying the metabolism of isolated mitochondria is understanding that *in vivo* mitochondria oxidize not just a single substrate but a mixture of metabolites coming from various metabolic pathways (Figure 1).

**Figure 1.**
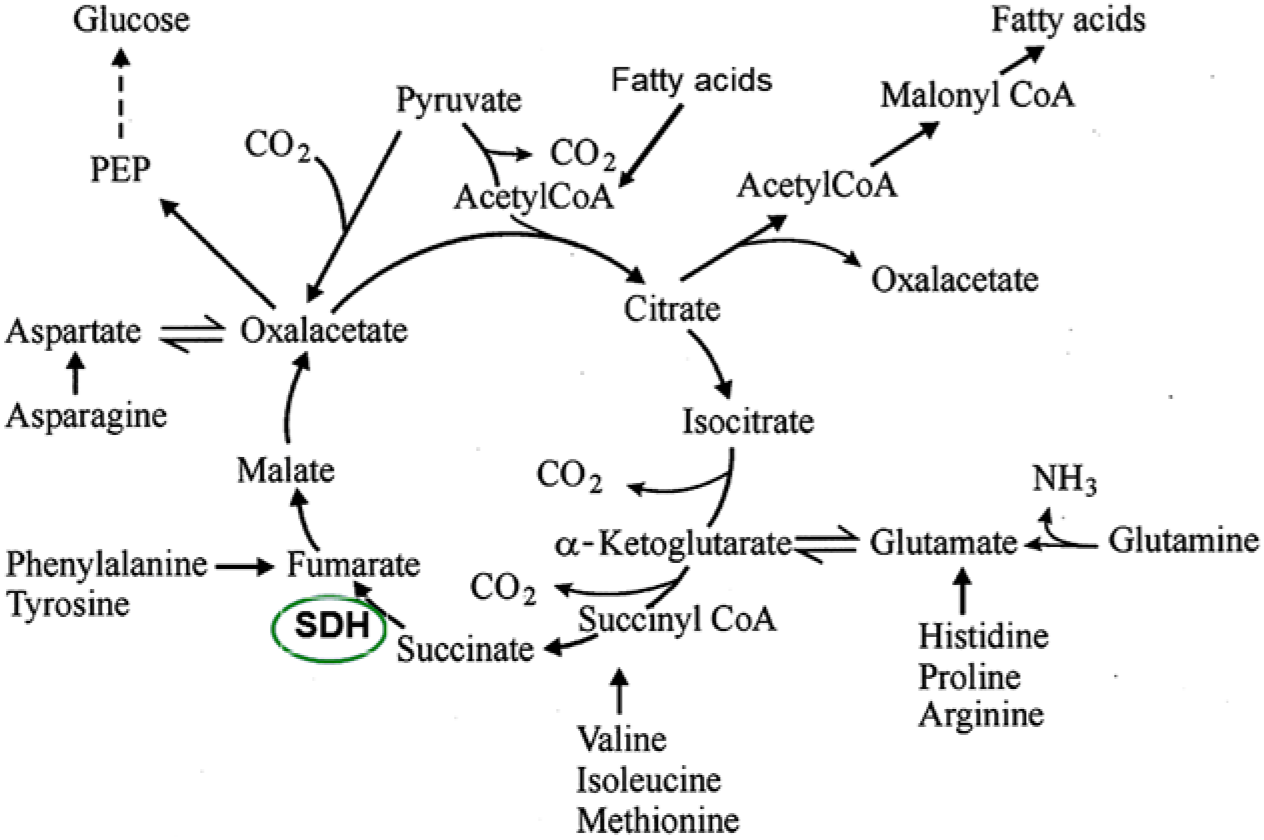
Metabolic interactions of the mitochondria tricarboxylic acids cycle with various metabolic pathways. The figure was adapted from [15].

Using mixtures of substrates, we have previously established that brain and heart mitochondria possess a potent intrinsic inhibition of succinate dehydrogenase (also known as the respiratory complex II), which is a mechanism of preventing the production of reactive oxygen species (ROS) when the organ is at rest (brain) or decreased functional activity (heart) [14,15]. We also established that substrate specificities and respiratory activities vary significantly between the organs and species in mitochondria. We have observed that a change in mitochondrial metabolic phenotype in transgenic animals resulted in the loss of the ability of the mutated *sod1* gene to cause symptoms of amyotrophic lateral sclerosis [16]. Therefore, to understand the mitochondrial role in maintaining the organ functions under normal and pathological situations, it is necessary to study the isolated mitochondria under metabolic conditions close to those *in vivo*.

## Materials and Methods

### Animals

Experiments to study rat heart and brain mitochondria were complied with the National Institutes of Health guidelines and were approved by the IACUC of the Carolinas Medical Center. Male Sprague Dawley rats (180–250 g) from Taconic Farms Inc. (Germantown, NY 12526) were used to isolate brain and heart mitochondria. All experimental procedures with mice C57Bl/6N were approved by the Vanderbilt and Mercer Institutional Animal Care and Use Committees.

### Isolation of mitochondria

Brain mitochondria were isolated from pooled forebrains of three rats using a modified method of Sims [17], as described in [18]. Rat heart mitochondria were isolated as described in [18]. For the isolation of kidney mitochondria, two kidneys from 1 C57Bl/6N mouse were cleaned from fat, cut into small pieces, and subjected to disintegration with polytron with two pulses for 2 seconds. Further procedures were performed similarly to the isolation of the heart mitochondria [18]. For the kidney mitochondria, we have found that with the final centrifugation at 8000g, the resulting mitochondria were of the same quality as mitochondria sedimented at 10000g followed by purification with the discontinuous Percoll gradient. This paper presents data for the kidney mitochondria isolated without purification with Percoll.

The isolation medium contained 75 mM mannitol, 150 mM sucrose, 20 mM MOPS, pH 7.2, 1 mM EGTA. The final suspensions of mitochondria were prepared using the incubation medium described below. Mitochondrial protein was determined with the Pierce Coomassie protein assay reagent kit.

### Measurements of mitochondrial respiration

We used different instrumentations for respiration measurements because the experiments with the rat brain, heart mitochondria, and kidney mitochondria were performed at different institutions and times. The respiratory activities of the rat heart and brain mitochondria were measured using a custom-made plastic minichamber of 560 μL volume equipped with a standard YSI (Yellow Spring Instrument Co., Inc.) oxygen minielectrode connected to a YSI Model 5300 Biological Oxygen Monitor [12,14]. The respiration rates of kidney mitochondria were measured using Fluorescence Lifetime Micro Oxygen Monitoring System (Instech Laboratories, Inc.) [19].

The incubation medium contained: 125 mM KCl, 10 mM MOPS, pH 7.2, 2 mM MgCl_2_, 2 mM KH_2_PO_4_, 10 mM NaCl, 1 mM EGTA, 0.7 mM CaCl_2_. At a Ca^2+^/EGTA ratio of 0.7, the free [Ca^2+^] is close to 1 μM as determined using Fura-2. The substrate concentrations were: 5 mM succinate without rotenone, 5 mM glutamate, 2.5 mM pyruvate, 2 mM malate. Oxidative phosphory;ation was initiated by addition of 150 μM ADP. L-palmitoyl-carnitine 0.025 mM dissolved in 50% ethanol. Substrates and substrate mixtures were added before the addition of 0.3 mg mitochondria. The data are presented as mean ± standard error of 3-5 separate experiments for each substrate and substrate mixtures.

### Chemicals

Chemicals were of the highest purity available. All solutions were made using the glass bidistilled water.

### Statistics

Comparisons between two groups were made by unpaired *t*-test.

## Results

### Endogenous substrates

The freshly isolated mitochondria from various organs may have different amounts of endogenous substrates in the mitochondrial matrix. Extreme examples are mitochondria from the liver, which can respire for as long as 15 minutes without added substrates, whereas isolated synaptic mitochondria, commonly referred to as the brain mitochondria, have zero oxygen consumption.

The contents of endogenous substrates in mitochondria from the isolated heart and kidney varied between these two extremes (not shown), but the kidney mitochondria are closer to the brain’s ones. However, even with added substrates, the composition and the amount of endogenous mitochondrial metabolites may affect respiration. For example: with the heart mitochondria, the oxidation rates of palmitoyl-carnitine + malate strongly depend on the presence of mitochondrial metabolites that serve as supporting substrates. Thus, the respiration rates depend on whether an experiment is performed shortly after the isolation or later [12,15].

In the mouse kidney mitochondria, with pyruvate alone, as an added substrate, the State 4 and State 3 oxygen consumption rates are roughly significantly slower than with added pyruvate + malate (Figure 2).

**Figure 2.**
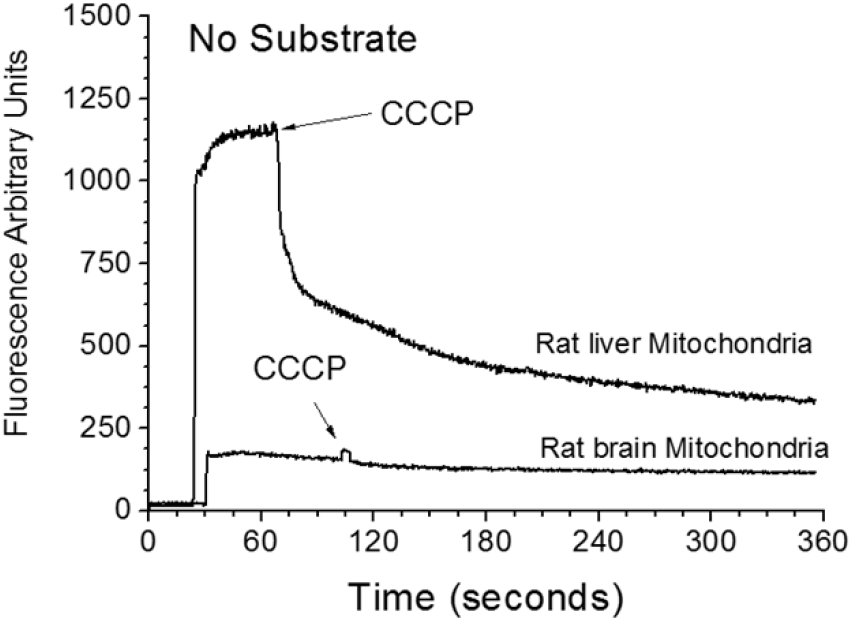
The mitochondrial NAD(P)H fluorescence in the freshly isolated rat liver and brain mitochondria incubated without added substrates. The figure was adapted from [13].

With glutamate as the sole substrate, the rate of the State 4 respiration was inhibited only slightly, compared with respiration with glutamate 5 mM+malate 2 mM, but the phosphorylating respiration (State 3) was (not shown). Unfortunately, we did not conduct similar experiments with the brain and heart mitochondria, which we performed some years ago, and at that time, we always added to mitochondria malate together with pyruvate and glutamate.

In the case of kidney mitochondria, we observed that isolated mitochondria contain few endogenous substrates. For this reason, oxidation of pyruvate requires malate as a source of oxaloacetate for entering the tricarboxylic acid (TCA) cycle. With glutamate as a substrate, malate provides oxaloacetate for the transamination reactions.

### Intrinsic inhibition of succinate dehydrogenase (SDH)

From the early days of mitochondriology, it was known that isolated mitochondria from the heart and brain very poorly oxidized succinate. SDH inhibition was not observed when mitochondria were isolated in the presence of defatted bovine serum albumin (BSA) [20]. For these reasons, the intrinsic inhibition of SDH was regarded for decades as an artifact of the isolation procedure. However, we have established that the intrinsic inhibition of SDH is a crucial physiological mechanism for preventing excessive production of ROS under conditions of diminished functional activity of the heart or brain [20].

Figure 4A shows no inhibition of succinate oxidation in the resting (State 4) and phosphorylating (State 3) respiration in the freshly isolated kidney mitochondria compared with the isolated brain (Fig.3B) and heart (Fig.3C) mitochondria.

**Figures 3.**
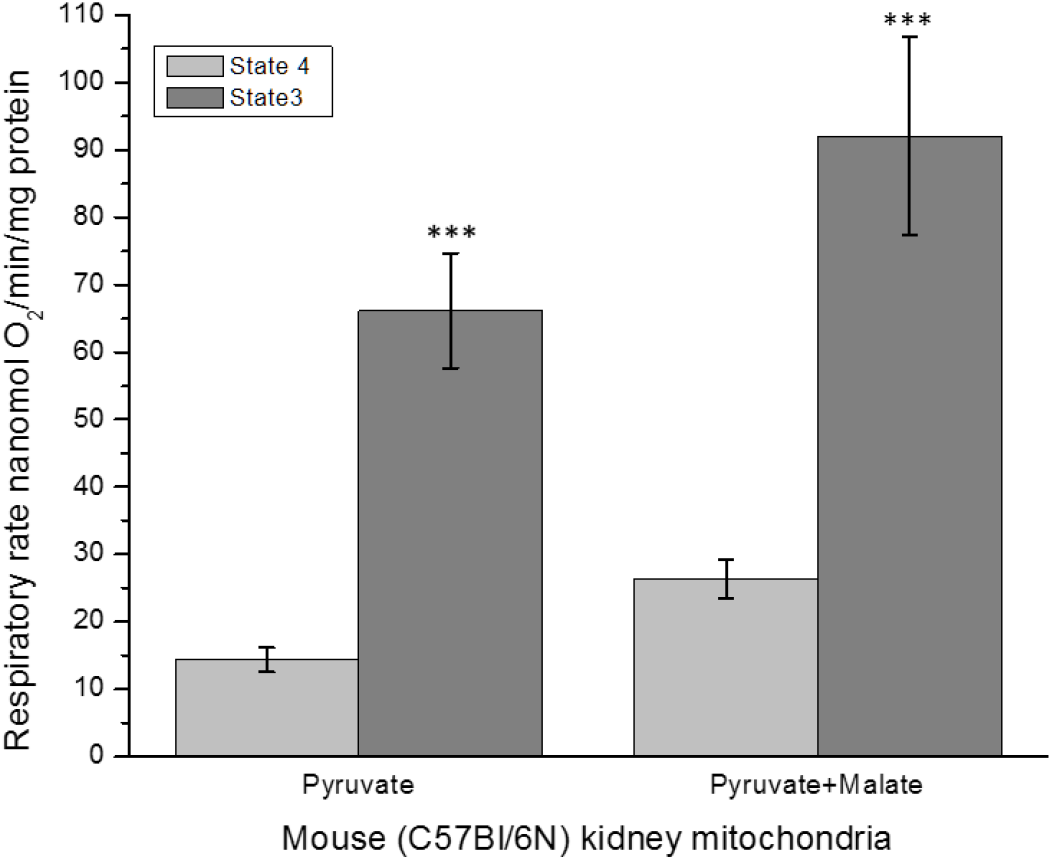
Effect of malate on oxidation of pyruvate by mouse kidney mitochondria. Oxygen consumption rates were measured without ADP (State 4) or with ADP supplementation (State 3). The data are mean ± standard error. *P<0.001 State 3 vs State 4 (n=5).

**Figure 4.**
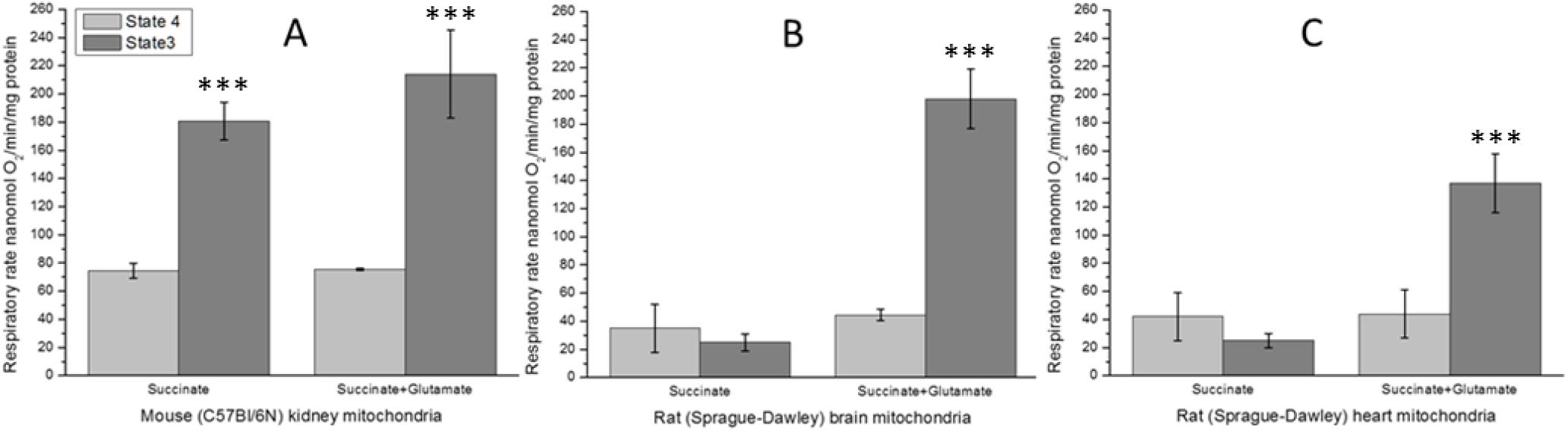
Kidney mitochondria do not possess the intrinsic inhibition of succinate dehydrogenase. Oxygen consumption was measured without ADP (State 4) or with ADP supplementation (State 3). The data are mean ± standard error. ***P<0.001 vs State 4 (n=5).

Isolated mitochondria from the brain and heart oxidize succinate poorly in the absence of rotenone, particularly in the presence of ADP. The addition of glutamate releases SDH inhibition by removing oxaloacetate from the active center in the transaminase reaction [13].

## Oxidation of L-palmitoyl-carnitine

Physiologists a long time ago have shown that heart and kidney mitochondria utilize long-chain fatty acids as the primary source of energy [8,11]. However, researchers at the mitochondria level rarely utilized long-chain fatty acids, such as palmitoyl-carnitine, as substrates for mitochondrial energization due to experimental challenges.

Figures 5 and 6 show that kidney, brain, and heart mitochondria have low respiration rates in States 4 and 3, with palmitoyl-carnitine + malate as the only substrates. Uncoupling led to severe inhibition of oxygen consumption (not shown). However, when other mitochondrial metabolites such as succinate, pyruvate, or glutamate have been added together with palmitoyl-carnitine, the rates of oxidative phosphorylation increased several-fold, and uncoupled respiration was high and linear (not shown, but see [12,13]). We named these mitochondrial substrates as supporting substrates for the oxidation of the long-chain fatty acids [12, 13].

**Figure 5.**
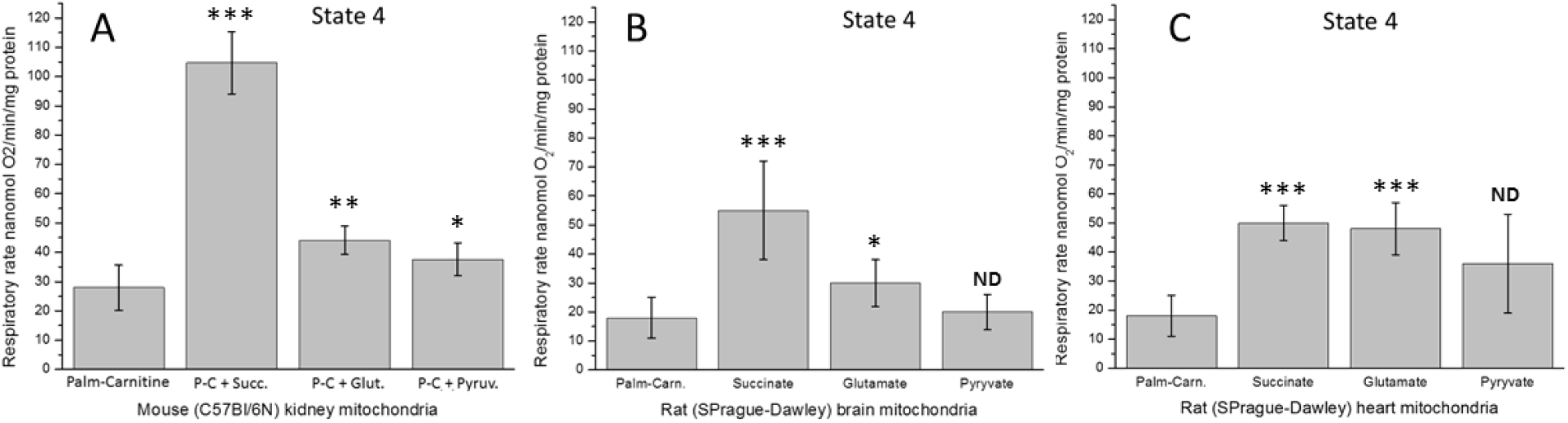
Effects of succinate, glutamate, and pyruvate on the resting oxidation of palmitoylcarnitine by the isolated kidney, brain, and heart mitochondria. Graphs show the oxygen consumption rates without ADP (State 4). The data are mean ± standard error. ***P<0.001 Palm-Carnitine vs P-C+Succinate (n=5); **P<0.01 Palm-Carnitine vs P-C+Glutamate; *P<0.05 and ND Palm-Carnitine vs P-C+Pyruvate

**Figure 6.**
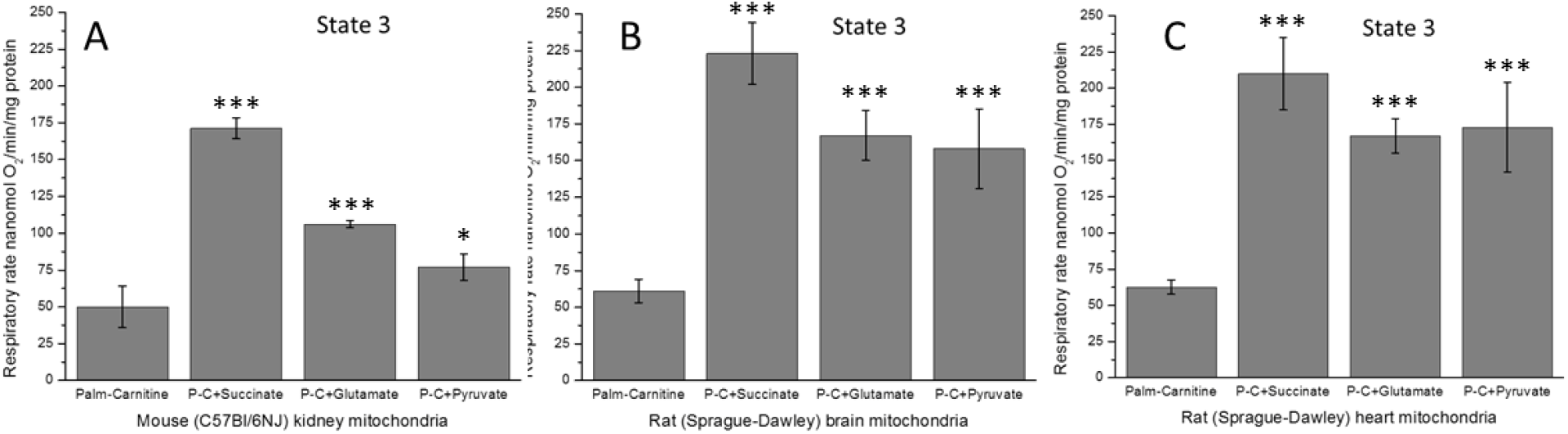
Effects of succinate, glutamate, and pyruvate on the rates of oxidative phosphorylation during oxidation of palmitoyl-carnitine by the isolated kidney, brain, and heart mitochondria. Graphs show the oxygen consumption rates with ADP (State 3). The data are mean ± standard error. ***P<0.01 Palm-Carnitine vs P-C+Succinate, P-C+Glutamate, and P-C+Pyruvate; *P<0.05 Palm-Carnitine vs P-C+Pyruvate (A) (n=5).

### Effects of supporting substrates on oxidation of palmitoyl-carnitine during resting State 4

Resting respiration (State 4) is the respiration of non-phosphorylating mitochondria. We observe State 4 respiration before adding ADP and after mitochondria phosphorylate the added ADP in experiments. In the Percoll purified mitochondria from the brain, kidney, and heart, both State 4 respiration rates are almost the same, indicating mitochondrial structural integrity.

The Discussion section provides a more detailed deliberation of respiration rates in State 4. Here we show that compared to the State 4 oxygen consumption rates by mitochondria oxidizing palmitoyl-carnitine + malate, taken as 100%, the presence of supporting substrates led to a severalfold increase in the rates of respiration in the kidney, brain, and heart mitochondria. The increases depended on the type of the supporting substrate and the origin of mitochondria. Succinate caused the most significant increases of the State 4 respiratory rates: mouse kidney mitochondria (MKM) 360%, rat brain mitochondria (RBM) 300%, and rat heart mitochondria (RHM) 280%. With 5 mM glutamate, the corresponding results were: MKM 160%, RBM 170%, and RHM 270%, and with 1.5 mM pyruvate, the results were: MKM 140%, RBM 110%, and RHM 200%.

### Effects of supporting substrates on oxidation of palmitoyl-carnitine in the presence of ADP (State 3)

The data presented in Table 1 demonstrate significant distinctions among supporting substrates in stimulating the phosphorylating oxidation of palmitoyl-carnitine. Succinate was the most compelling supporting substrate with the kidney, brain, and heart mitochondria.

In comparison with the respiration rate with P-C + malate as substrates, taken as 100%, the addition of 5 mM succinate increased the rate of the State 3 respiration in the MKM 340%, RBM 370%, and the RHM 340%. The corresponding effects of 5 mM glutamate were: MKM 200%, RBM 270%, and RHM 270%. The effects of 1.5 mM pyruvate on the P-C + malate State 3 oxidation were: MKM 150%, RBM 260%, and RHM 280%. A recent publication by Berezhnov et al. [21] showed the increase in State 3 respiration and energization upon the addition of 10μM palmitoyl-carnitine to the heart mitochondria oxidizing malate+α-ketoglutarate (Fig. 5A) or glutamate+pyruvate (Fig, 5B,D).

## Discussion

This work shows very distinct metabolic properties of murine kidney mitochondria compared with the brain and heart mitochondria. First, kidney mitochondria are highly efficient in succinate utilization, and they do not encounter the potent intrinsic inhibition of SDH observed in the brain and heart mitochondria. Second, similar to the brain and heart mitochondria, kidney mitochondria also increase β-oxidation of long-chain fatty acids in the presence of supporting substrates, but quantitatively the stimulation may be different. Our data align with the previously reported physiological reliance of kidney energy metabolism on fatty acid β-oxidation. Furthermore, our findings highlight the critical role of succinate and long-chain fatty acids in kidney energy metabolism. We recently showed that 58% of glutamate-driven respiration was complex II-dependent [19]. Kidney mitochondria adapt to the high energy demands by diverting the mitochondrial metabolite flux to succinate to feed the Complex II-mediated respiration. Succinate is produced via transamination of pyruvate or glutamate to accelerate the rates of mitochondrial respiration and ATP production.

Furthermore, Complex II has a higher abundance than Complex I, and Complex I activity is susceptible to oxidative stress. We suggest that the presence of succinate during LCFAs β-oxidation is also necessary for accelerating the redirection of electrons from CoQH2 into the respiratory chain. We suggest that increased reliance on complex II and fatty acid β-oxidation supports renal mitochondria’s metabolic plasticity.

### The main reasons why long-chain fatty acids are the primary substrates for the hardworking organs

Natural long-chain fatty acids (LCFAs) have an even number of carbons, from 14 to 20, that can be saturated or unsaturated. Polyunsaturated fatty acids with three or more double bonds are usually not utilized as a source of energy. However, they are present in phospholipids that construct the biological membranes and serve as substrates to produce the signaling molecules, such as prostaglandins [22,23]. The reserves of fatty acids in triglycerides are vast, measured in kilograms.

Unlike the LCFAs, glucose storages are very limited, and glucose is in great demand for the biosynthesis of many other vital molecules: nucleic acids, purine, and pyrimidine nucleotides, glycolipids and glycoproteins, etc. Therefore, the usage of glucose as a source of energy is somewhat limited. The absolute demands for glucose have only erythrocytes that consume at least 240-360 grams of glucose in a person of average size in a day (A. Panov, unpublished data). Even the central nerve system satisfies its energy demands by more than 20% for the expense of β-oxidation of LCFAs [reviewed in 13,24]. In kidneys, glucose has a non-energetic function of reabsorption sodium ions from the glomerular filtrate back to the circulation.

Amino acids have no storage in the body and are constantly formed during catabolism of the food or organism’s proteins and formed anaplerotically.

Another critical reason why fatty acids are essential as a source of energy is that glucose cannot provide fast rates of ATP production because the NADH dehydrogenase of the respiratory complex I limits the overall transport of electrons in the respiratory chain of mitochondria [25]. Glutamate and pyruvate cannot be regarded as the rigid substrates for the complex I because, in the brain, heart, and kidney, these substrates, to a large degree, undergo transamination with the formation of succinate, which has a much faster rate of oxidation and can drive the reverse electron transport [14,19].

According to Brandt, during β-oxidation of LCFAs, mitochondria reduce the NAD to NADH + H^+^ and the mitochondrial pool of ubiquinone to ubiquinol (CoQH_2_) [26]. In the mitochondria, oxidizing LCFAs, the electrons from the CoQH_2_ at the respiratory complex II (SDH) level become redirected into the electron transport chain, where the respiratory complex III instantly carries electrons to complex IV reducing O_2_ to H_2_O [26–28]. In the well-energized resting mitochondria (State 4 respiration), the excess of electrons is transferred to complex I and accelerates the production of superoxide radicals [12,13].

Thus, in the organs consuming a large amount of ATP, only β-oxidation of LCFAs can maintain the high rates of oxidative phosphorylation for a long time.

### Reasons for low endogenous substrates content and lack of the intrinsic inhibition of SDH in kidney mitochondria

The kidney, brain, and heart are the fastest oxygen-consuming organs in the human body. In a human at rest, the oxygen consumption index (OCI), which is the ratio: % O_2_ consumption by the organ/ % of the total body weight, is correspondingly 12 for the kidneys, 10 for the brain, and 27.5 for the heart [29]. However, at the peak of functional activity, the brain increases the rate of O_2_ consumption 2-3 fold, and the heart may increase O_2_ consumption and energy production nine times [30], whereas kidneys work at all times almost at the same pace.

In order to satisfy high energy demands, kidney, heart, and brain mitochondria have the vast surface of the inner mitochondrial membrane and a large number of the respiratory chain complexes organized in the respirosome supercomplexes [31,32]. The matrix volume is small, particularly in comparison with the liver mitochondria, and the content of the matrix is a hard gel [33]. Brain mitochondria are typically relatively small, whereas kidney and particularly heart mitochondria are very large. Therefore they may have some osmotically active matrix volume and thus contain a small amount of metabolites, whereas the brain mitochondria do not.

Mitochondria constantly produce reactive oxygen species (ROS), predominantly as superoxide radicals and a small amount of hydrogen peroxide [26]. However, significant production occurs only in resting mitochondria or when the consumption of ATP diminishes, like in the brain during sleep or at low physical activity in the heart. The most significant ROS production occurs due to reverse electron transport initiated by oxidation of succinate or β-oxidation of FAs, which also involves SDH participation [26]. Therefore, as we have suggested earlier, the intrinsic inhibition of SDH represents the internal mechanism of preventing the excessive production of ROS at lower functional loads of the heart or brain [15]. For the first time, we show in this paper that in the kidneys that work constantly, the intrinsic inhibition of SDH is absent.

### Effect of supporting substrates on the resting respiration in mitochondria oxidizing palmitoyl carnitine

In the absence of oxidative phosphorylation, respiring mitochondria maintain maximal energization, and oxygen consumption rates depend on the intrinsic inner mitochondrial membrane proton and electrons leaks [34,35]. A significant proportion of the basal leak can be attributed to mitochondrial anion carriers, whereas the proton leak through the lipid bilayer appears minor [35]. Up to 30% of the proton leak of freshly prepared mitochondria can be prevented with bovine serum albumin (BSA) and was attributed to fatty acids [33]. In addition to proton leak, energized mitochondria also show electrons leak from the respiratory chain to the formation of superoxide radicals [35]. We have shown earlier that the significant increase in oxygen consumption during resting respiration of the heart and brain mitochondria in the presence of palmitoyl-carnitine and supporting substrates was accompanied by a manifold increase in ROS production [12,13]. The rise in oxygen consumption rates during resting respiration was probably the primary reason why fatty acids were considered as mitochondrial uncouplers for a long time. Classical uncouplers cause dissipation of the mitochondrial transmembrane potential as heat, thus diminishing the rate of ATP production. As is shown in Table 1 and earlier publications [12,13], oxidation of palmitoylcarnitine in the presence of supporting substrates results in several-fold increases in the rates of ATP production in the kidney, heart, and brain mitochondria. Thus increases in oxygen consumption during the State 4 respiration were caused not by the uncoupling, as was generally suggested earlier, but by the excessive provision of electrons and activation of ROS production due to the reverse electron transport, which occurs only in the highly energized mitochondria.

### Effect of supporting substrates on the phosphorylating respiration in mitochondria oxidizing palmitoyl carnitine

Figures 5 and 6 demonstrate that succinate provides, as a supporting substrate, the highest rates of oxidative phosphorylation with the kidney, brain, and heart mitochondria oxidizing palmitoylcarnitine. The exact mechanism of stimulating fatty acid oxidation by succinate, glutamate, and pyruvate remains elucidated. Based on the data presented by Brand and colleagues [26–28], we suggest, as a working hypothesis, that succinate stimulates the reverse transport of electrons from the membrane pool of CoQH_2_ into the respiratory chain. Our preliminary unpublished experiments have shown that the supporting effect of succinate was observed at the concentration of 0.5 mM. It is plausible that the supporting effects of glutamate and pyruvate may also be associated with the conversion of these substrates first to α-ketoglutarate and then to succinate via transamination reactions. Table I shows that pyruvate causes only slight stimulation of the State 3 oxygen consumption with kidney mitochondria, compared with succinate and glutamate, which can be explained by the particular role glucose plays in the reabsorption of Na^+^ ions in the kidney proximal tubules.

### Functional activities of the kidney tubules predetermine the mitochondrial substrate preferences

The kidneys in normoglycemic humans filter approximately 160-180 g of glucose per day, which is returned to the systemic circulation by the reabsorption in the proximal tubule [35]. The glucose reabsorption from glomerular ultrafiltrate has the significant function of the reabsorption of Na+ ions. Sodium-glucose cotransporters (SGLT) mediate apical sodium and glucose transport across cell membranes. Cotransport is driven by active sodium extrusion by the basolateral Na^+^/K^+^-ATPase, thus facilitating glucose uptake against the intracellular uphill gradient. Basolaterally, glucose exits the cell through facilitative glucose transporter 2 (GLUT2) [36].

In the kidney’s proximal tubule segment 1 and 2 (S1/2), SGLT2 accounts for more than 90% of glucose reabsorption from the glomerular ultrafiltrate, and SGLT1 is responsible for reabsorbing nearly 3% of the filtered glucose load in the proximal renal tubule segment 3 (S3) [reviewed in 37]. SGLT2 is positively linked to the Na^+^/H+-exchanger 3 (NHE3). Therefore, for SGLT2, the stoichiometry glucose/Na^+^ is 1, whereas, for the SGLT1, the stoichiometry glucose/Na^+^ is 2 [35]. The tight coupling of glucose and sodium reabsorption suggests that kidneys must avoid the utilization of glucose as a major source of energy. Moreover, during starvation kidney must provide enough glucose present in the ultrafiltrate for the active reabsorption of Na^+^ ions, which suggests that gluconeogenesis must occur in the podocytes.

### Functional and metabolic activities of glomerular podocytes

The glomerulus is a ball of capillaries surrounded by the Bowman’s capsule. Each glomerulus is composed of four types of cells: parietal epithelial cells (Bowman’s cells), visceral epithelial cells (podocytes), mesangial cells, and endothelial cells [38]. The glomerular podocytes have the cell body, cell- and foot processes. A detailed description of the structure and functions of glomerular podocytes has been published in papers and reviews, of which we mention only a few [38–40]. The podocytes contain numerous mitochondria to satisfy energy expenditures for the maintenance of the complex structure of the slit diaphragm and constant dynamic interactions with the basal membrane [38]. Although it is recognized that the primary source of ATP in podocytes is mitochondrial oxidative phosphorylation [38], some authors presented evidence on glycolytic production of ATP. However, these data cannot be regarded as decisive because the authors used antibiotics, which are mitotoxic, and nonphysiological substrates for the podocyte mitochondria [41,42].

The functional and metabolic roles of glomerular podocytes to a large degree can be compared with those of the astroglial cells in the brain. In astroglia, β-oxidation of LCFAs provides energy and carbons for aerobic production of lactic acid, which serves as a primary substrate for synaptic mitochondria and anaplerotic replenishment of glutamate [13,24]. We suggest that the glucose-dependent reabsorption of Na^+^ from the ultrafiltrate makes unlikely the possibility that kidney gluconeogenesis occurs in the tubule epithelial cells. Instead, glomerular podocytes have all conditions for maintaining active gluconeogenesis. The main gluconeogenic precursors in humans are lactate, glutamine, alanine, and glycerol account for 90% of overall gluconeogenesis [3,43]. Podocytes are surrounded by the plasma ultrafiltrate containing all these substrates. The tight structural organization of the β-oxidation enzymes and respirosome in mitochondria [43] with abundant fatty acids and supporting substrates maintains high energy and redox potentials in mitochondria, and the cytoplasm redirects pyruvate and oxaloacetate to the synthesis of glucose in podocytes. During starvation, glucose produced in the podocytes will be extruded to glomerular ultrafiltrate and thus participate in the reabsorptive co-transport of glucose and sodium to the central bloodstream.

## Conclusions

In the hard-working organs, the mitochondrial β-oxidation of the long-chain fatty acids in the presence of other mitochondrial metabolites is the primary energy source. The heart and brain mitochondria, which work in a wide range of functional loads, possess potent intrinsic inhibition of succinate dehydrogenase to prevent ROS production at low functional activities. Kidney mitochondria do not have intrinsic inhibition of SDH, probably because they constantly work hard at all times. The methodology to study β-oxidation of LCFAs in the presence of supporting substrates opens a new opportunity to study energy metabolism and metabolic phenotypes of mitochondria from different organs and species under normal and pathological conditions.

## Acknowledgments

This work was supported by funding from the National Institutes of Health (R01HL124116 and RO1HL157583) and Navicent Health Foundation grant (Mercer University School of Medicine).

